# RNA programmable cell targeting and manipulation with CellREADR

**DOI:** 10.1101/2024.11.26.625312

**Authors:** Xiaolu Yang, Kehali Woldemichael, John Hover, Shengli Zhao, Xiao Guo, Yongjun Qian, Jesse Gillis, Z. Josh Huang

## Abstract

Methods that provide specific, easy, and scalable experimental access to animal cell types and cell states will have broad applications in biology and medicine. CellREADR - Cell access through RNA sensing by Endogenous ADAR (adenosine deaminase acting on RNA), is a programmable RNA sensor-actuator technology that couples the detection of a cell-defining RNA to the translation of an effector protein to monitor and manipulate the cell. The CellREADR RNA device consists of a 5’ sensor region complementary to a cellular RNA and a 3’ payload coding region; payload translation is gated by the removal of a STOP codon in the sensor region upon base pairing with the cognate cellular RNA through an ADAR-mediated A-to-I editing mechanism ubiquitous to metazoan cells. CellREADR thus highlights the potential for RNA-based monitoring and manipulation of animal cells in ways that are simple, versatile, and generalizable across tissues and species. Here, we describe a detailed protocol for implementing CellREADR experiments in cell cultures and in animals. The procedure includes sensor and payload design, cloning, validation and characterization in mammalian cell cultures. The in vivo animal protocol focuses on AAV-based delivery of CellREADR using brain tissue as examples. We describe current best practices, various experimental controls and trouble-shooting steps. Beginning from sensor design, the validation of RNA sensor-actuators in cell cultures can be completed in 2-3 weeks. The construction of AAV vectors and their in vivo validation and characterization will take additional 6-8 weeks.

## Introduction

The diversity of cell types and cell states underlies tissue organization and systems function within individual organisms and across species. Cell types and states are fundamentally shaped by their gene expression profiles. Advances in single-cell RNA sequencing is accelerating the identification of many if not all molecularly-defined cell types in humans and in many other organisms^1–3^. Beyond cell type discovery and cataloging, it is necessary to systematically monitor and manipulate each and every cell type in order to identify their specific roles in tissue organization and system function. Cell-type-specific monitoring and intervention is also key to precision diagnosis and therapeutics in medicine. Achieving these goals requires cell type technologies that are specific, easy, scalable, and generalizable across tissues and species. We recently invented CellREADR - Cell access through RNA sensing by Endogenous ADAR (adenosine deaminase acting on RNA), a programmable RNA sensor-actuator technology that couples the detection of a cell-defining RNA to the translation of an effector protein to monitor and manipulate the cell^4^. We further show that CellREADR can perform logical AND and OR gate operations in RNA sensing. Essentially the same method is developed by two other groups and termed RADARS (reprogrammable ADAR sensor)^5^ or RADAR (RNA sensing using adenosine deaminases acting on RNA)^6^, respectively. Importantly, we further demonstrate that CellREADR enables recording and manipulating cell types in behaving animals and in human ex vivo tissues^4^. Together, these studies highlight the potential of this technology for RNA-based monitoring and editing of animal cells in ways that are versatile, scalable, programmable, and generalizable across organ systems and animal species.

At its core, CellREADR harnesses an RNA sensing and editing mechanism ubiquitous to metazoan cells to detect the presence of specific cellular RNAs and gate the translation of effector proteins to monitor and manipulate the cell (**Figure 1**). RNA editing is a widespread and robust post-transcriptional mechanism essential to animal cells and is implicated in recoding, splicing regulation, microRNA targeting, cellular innate immunity, and other RNA modulatory processes^7,8^. The most prevalent form of RNA editing is adenosine-to-inosine (A→I) conversion catalyzed by ADARs; inosine is recognized as guanosine (G) by the cellular machinery^8^. ADARs recognize and are recruited by stretches of base-paired double-stranded RNAs^8^, and therefore can operate as a sequence-guided base editing machine. The mammalian ADAR gene family is composed of several orthologs and isoforms with different tissue, cellular, and subcellular expression patterns and catalytic activities^7–9^ We design CellREADR as a single modular readrRNA molecule, which consists of a 5’ sensor domain of ∼250 nucleotides complementary to a cellular target RNA and an in-frame 3’ domain that encodes a payload protein; the sensor domain further contains an ADAR-editable STOP codon that acts as a conditional translation switch^4^ (**Figure 1**). The entire readrRNA is several kilobases in length, largely depending on the size of the effector included. CellREADR can be delivered to animal cells through DNA expression plasmids as well as *in vitro* synthesized RNAs. In cells expressing the target RNA, the sensor domain of readrRNA forms a double-strand RNA with the target, which recruits ADAR to convert the UAG STOP codon to a UGG tryptophan codon, thereby switching on payload translation.

**Figure 1.**
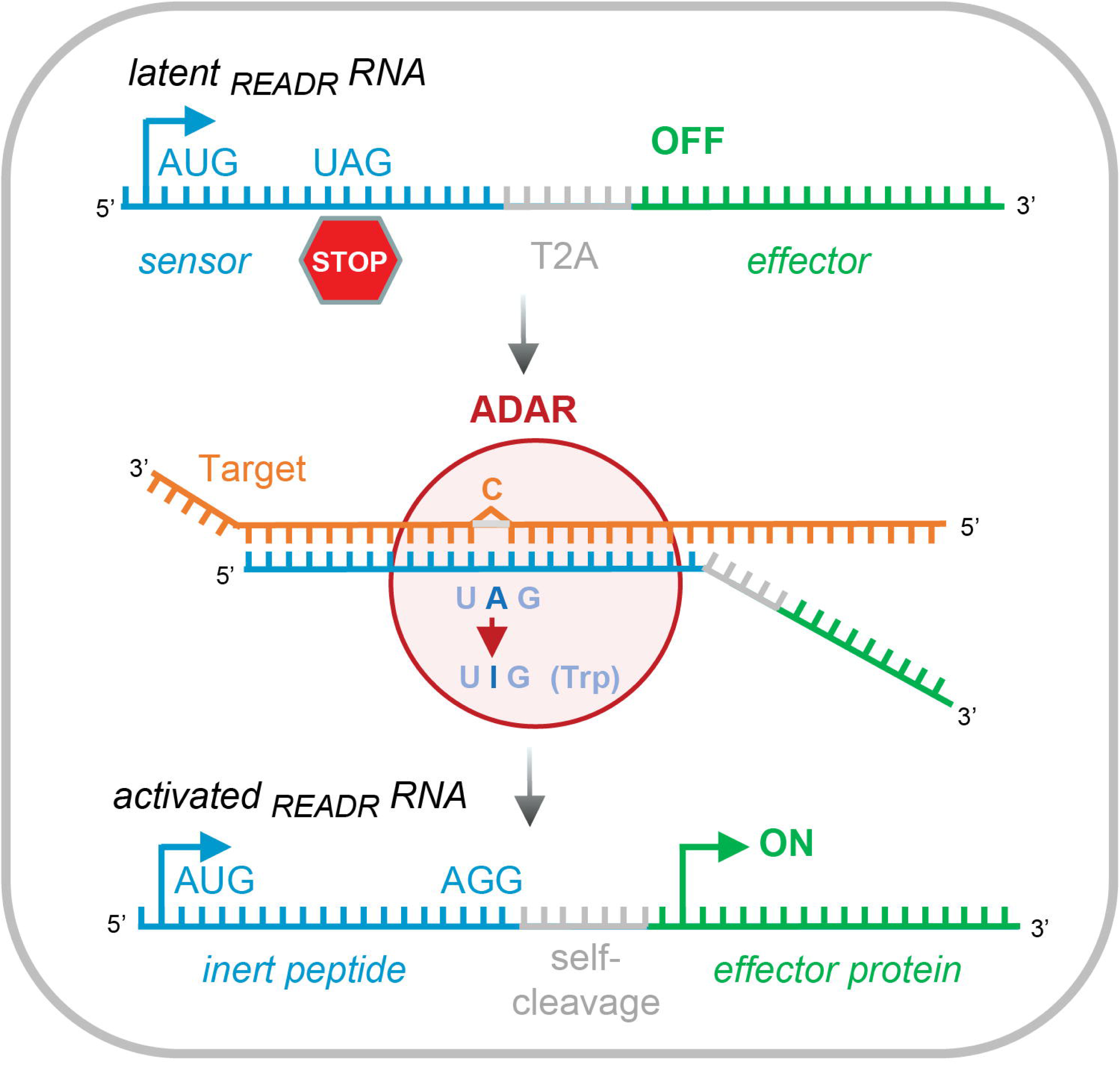
Schematic of CellREADR design. CellREADR is designed as a single modular readrRNA molecule, consisting of a 5′ sensor domain and 3′ effector domain. The sensor domain is ∼250 nucleotides long and complementary to a cellular target RNA. An editable STOP codon is designed in-frame around the center of the sensor domain to gate the translation of the effector protein. Downstream of the sensor domain is a sequence encoding the self-cleaving peptide T2A, followed in-frame by the 3’ effector domain. Base-pairing between the sensor domain and target RNA sequence recruits the ADAR protein to assemble an editing complex. ADAR converts the UAG STOP codon to UIG, which is recognized by the cellular machinery as UGG tryptophan codon, thereby switching on the translation of the effector.

The defining features of CellREADR are that it operates as a single RNA molecule, leverages sequence base pairing for cellular RNA sensing, and actuates payload translation through a robust endogenous RNA editing mechanism ubiquitous to animal cells. In essence, CellREADR reduces the cell type/state access problem in biology and medicine to a cellular RNA sequence detection and actuation problem. As such CellREADR represents a next-generation technology that is inherently 1) specific to cells defined by RNA markers; 2) easy to build, use, and disseminate; 3) scalable for targeting RNA-defined cells across tissues; 4) generalizable across animal species including human; and 5) programmable to achieve intersectional targeting of cells defined by two or more RNAs as well as multiplexed targeting of several cell types in the same tissue.

DART VADAR (Detection and Amplification of RNA Triggers via ADAR) is a variation of CellREADR, which features a payload design that amplifies the signal from STOP codon editing in the sensor domain^10^. This amplification is mediated by adding a hyperactive and minimal ADAR variant to the payload coding cassette to facilitate STOP editing in the sensor domain, thereby constituting a positive feedback loop. This design may further improve the dynamic range of CellREADR.

### Delivery

CellREADR can be delivered to cells either through DNA expression vectors or as in vitro synthesized RNAs. These two forms of delivery differ substantially in readrRNA production, subcellular localization, half-life, chemical modifications, and thus CellREADR functionality and applications **(Figure 2).**

**Figure 2.**
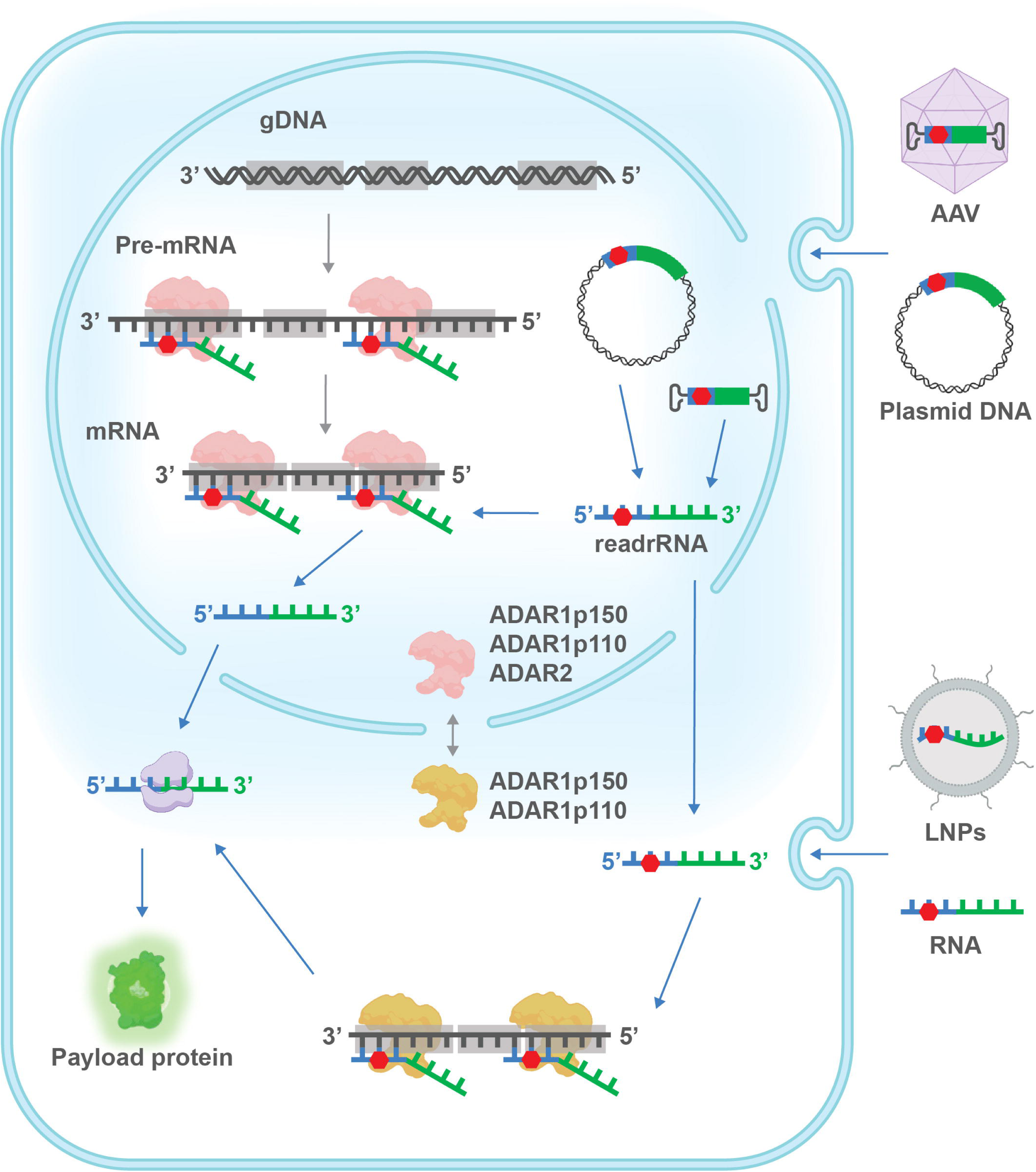
Two modes of CellREADR delivery and action. CellREADR can be delivered to cells either as DNA vectors or as in vitro synthesized RNAs. DNA expression vectors can be viral vectors and non-viral vectors. Viral vectors can be delivered through viral transduction, including AAVs, adenovirus and lentivirus. Non-viral vectors can be delivered by injection, electroporation and liposome. Latent readrRNAs are transcribed in the nucleus and then base pair with cellular target pre-mRNA and mRNA. Nucleus localized ADAR proteins are recruited and catalyze the A-to-I editing on the mismatched Adenosine. Latent readrRNAs are thus activated and exported to cytoplasm, as unedited readrRNAs are also exported to cytoplasm. Exported latent readrRNAs can base pair with cytoplasmic target mRNAs and be edited by cytoplasmic localized ADARs. ADAR2 is considered be localized to the nucleus, while both isoforms of ADAR1 shuttle between nucleus and cytoplasm. Activated readrRNAs in the cytoplasm are translated to payload proteins. Alternatively, in vitro synthesized readrRNAs can be packaged into lipid nanoparticles or viral like particles. After being delivered to cells through endocytosis, latent readrRNAs are largely restricted to the cytoplasm. They mainly base pair with the target mRNA and are edited by ADAR1p150 and ADAR1p110 to trigger payload translation.

#### DNA expression vectors

DNA vectors can be delivered using viral vectors (e.g. AAV, lentivirus), transfection (including electroporation), or nuclei injection. With this delivery route, readrRNAs are co-transcribed with the cellular target RNA in the nucleus. RNA sensing and editing likely can occur soon after transcription and during target pre-mRNA splicing. Indeed, both ADAR1 (p110) and ADAR2 isoforms are abundant and enzymatically active in the nucleus^7–9^. RNA sensing and editing likely continue after nuclear export of readrRNA and target mRNAs to cytoplasm, as the two ADAR1 isoforms p110 and p150 shuttle between the nucleus and cytoplasm^7–9^ Therefore, continued co-production of readrRNA and target RNA will engage the full RNA life cycle for RNA sensing and editing, with long-lasting payload expression (**Figure 2**).

#### In vitro synthesized RNAs

Similar to mRNA vaccines^11,12^, readrRNAs can be generated by in vitro transcription, chemically modified where necessary, packaged into lipid nanoparticles or viral like particles, and delivered to cells through endocytosis. With this delivery route, readrRNAs are expected to be largely restricted to the cytoplasm and thus only or mainly sense the target mRNA. readrRNAs likely have a half-life of on the order of tens of hours, and mediate payload translation during this time window **(Figure 2)**. This more transient modality of CellREADR function can be either an advantage or disadvantage, depending on the application, especially in the context of therapeutics.

### Applications

As a “general purpose” RNA sensor-actuator device built upon an endogenous RNA editing mechanism ubiquitous to metazoan cells, CellREADR has diverse applications in basic biological research and potentially in RNA therapeutics. CellREADR can sense RNAs that define living cell states to gate the translation of any payload protein of interest, and thus to label, record, and manipulate the cell according to research or therapeutic need. This wide range of applications can be implemented in many tissues and animal species, as long as there is a method to deliver DNA or RNA to the intended cells. In addition, many RNAs are generated with multiple isoforms, such as splice variants, alternative promoters or 3’UTRs, and circular RNAs^13–15^, in different tissues and cells of the organism, but the biological significance of these isoforms is often unclear. CellREADR can potentially detect these RNA isoforms in their resident living cells and reveal the significance of these alternative transcripts in the context of cell physiology and tissue organization and function.

### Comparison with other methods

Most genetic approaches to accessing cell types rely on DNA-based transcriptional mechanisms to mimic certain cell-specific RNA expression (**Figure 3**), often through germline engineering.

**Figure 3.**
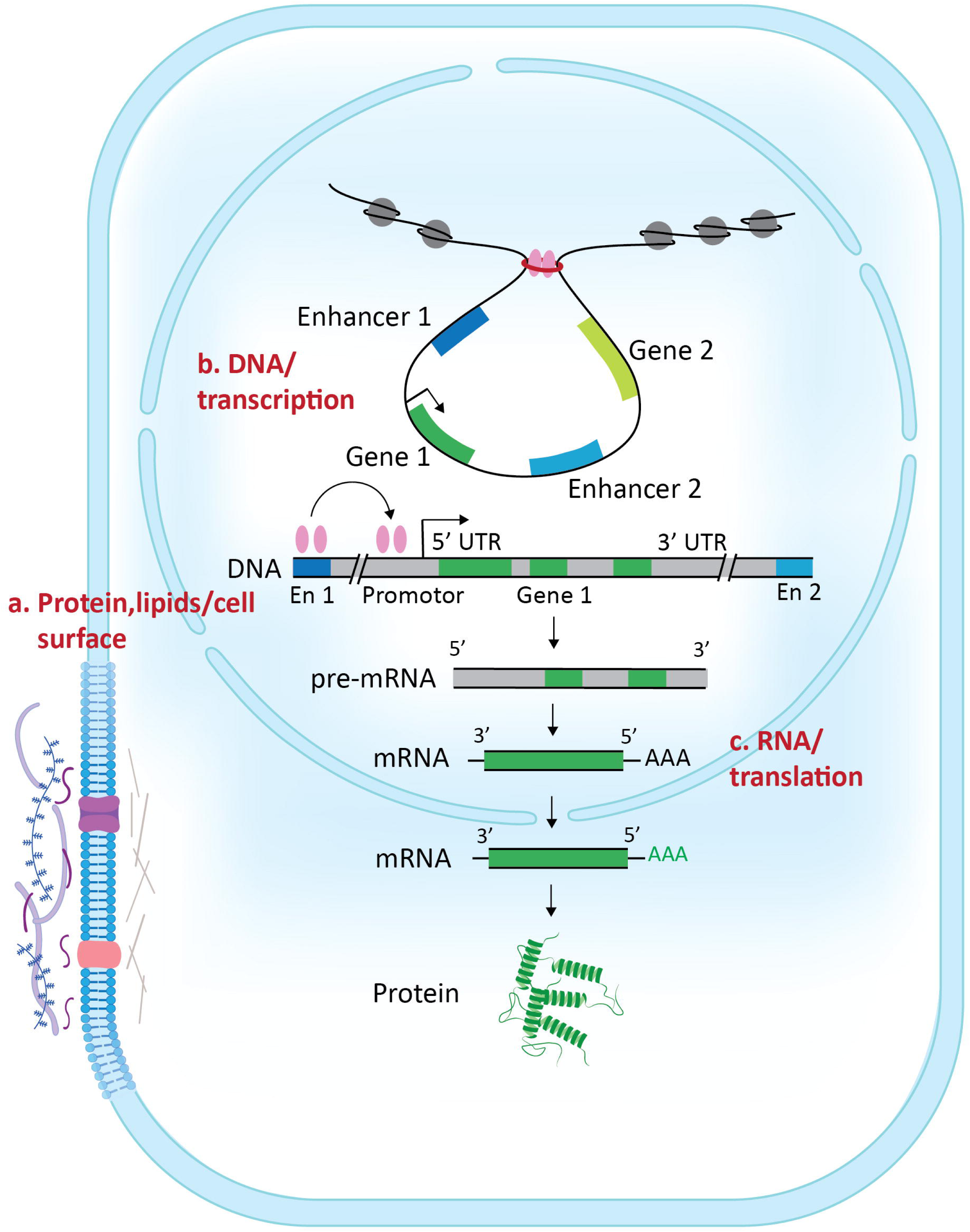
Strategies for cell type access. A general scheme of cellular gene expression illustrating the three major strategies for gaining cell type access. a) Viral capsid engineering or lipid nanoparticles engage cell surface proteins or lipids to enter the cell. b) Cell type-specific transcriptional enhancers direct payload transcription through either germline engineering or viral vectors. c) RNA sensors (e.g. CellREADR) can be delivered by AAV or LNP to detect specific cell-defining RNAs and gate payload translation.

However, all germline approaches are inherently cumbersome, difficult to scale and generalize across species, and raise ethical issues especially in primates and humans^16,17^. Recently, transcriptional enhancer-based viral vectors have shown promise for targeting cell types^18–20^. Indeed, large scale epigenome and transcriptome analyses in multiple mammalian species have identified tens of thousands of putative enhancers. However, enhancers operate through DNA-protein interactions with nuanced relationships to target genes and cell types^21^. Thus cell type enhancers are difficult to identify and validate, often require large-scale efforts beyond the capability of most academic labs^18^. Furthermore, as most enhancers are first identified in a few model organisms (e.g. mice), their transferability to other species, especially evolutionarily more distant species, remains uncertain. Therefore, whether the enhancer approach can truly scale and generalize across species, especially those without a high-quality genome assembly, remains to be demonstrated.

As cell types are often defined by RNA profiles, an alternative and more direct approach compared with DNA and transcription-based method is to control the expression of payload genes based on the presence of certain cellular RNAs (**Figure 3**). For example, one group designed a splicing-based approach for controlling transgene expression whereby separate translational reading frames are coupled to cell-specific alternative exons^22^; however, splicing patterns do not generally distinguish most cell types and states. Another group described a Ribozyme-Enabled Detection of RNA method (RENDR), which uses cellular transcripts to template the assembly of split ribozymes, triggering splicing reactions that generate orthogonal protein outputs in bacteria^23^. In addition, the toehold switch is a strategy to control translation by the detection of a cellular RNA, also initially developed for bacterial cells^24,25^; but this method is more difficult to implement and has limited dynamic range in eukaryotic cells^25,26^. In this context, CellREADR represents a simple and generalizable RNA-based approach to cell types and cell states across metazoan species.

In addition to leveraging gene expression, a categorically different approach to cell types is to engage cell membrane and surface properties (e.g. receptors, extracellular matrix) that can be distinguished by delivery vehicles such as viral particles or lipid nanoparticles (**Figure 3**). Indeed, major efforts and progress have been made in the past decade to engineer and evolve recombinant AAV capsids for delivering gene expression vectors to diverse tissues and species^27–29^. In addition, lipid nanoparticles (LNPs) are formulated and conjugated to surface recognition proteins (e.g. antibodies) to deliver RNAs to mostly mammalian tissues and cell types^30,31^. However, while engineered AAV capsids and LNPs can bias to or against payload delivery to certain tissues, these cell surface-based methods are fundamentally limited in accessing finer resolution cell types and cell states within tissues. On the other hand, proper delivery systems are prerequisite for all gene expression-based approaches to cell targeting: cells resistant to delivery vehicles preclude the application of gene expression-based approaches, and favorable tissue bias in delivery vehicles facilitate and synergized with gene expression-based approaches. This point is discussed in more detail for in vivo animal applications of CellREADR.

### Current limitations

The current version of CellREADR can be improved in specificity, sensitivity, efficacy (i.e. payload expression level), and programmability.

#### Specificity

Reliable design of specific sensor sequence is key to CellREADR application. Most target RNAs contain many CCA sequences as potential “anchor sites” for designing editable T<u>A</u>G STOP in the sensor region. In the target pre-mRNA, these CCA sites are distributed across exons, introns, 5’ and 3’UTRs; in the target mRNA, CCA sites are distributed across coding regions (CDS) and UTRs. Currently, we do not yet understand the rules or principles of sensor design. In practice, we often design approximately a dozen sensors to test in cell cultures. As most target RNAs that define cell type of interest are not expressed in commonly used cell lines (e.g. human HEK293, mouse N2A), we need to exogenously express these targets in the cell line. However, RNA sensors validated in cell cultures may not always translate to the intended cell type in animal tissues in vivo; currently, approximately 50% cell culture-tested sensors perform well in animals^4^.

It is possible that differences in RNA content, ADAR isoforms and expression levels may reduce the correspondence of sensor performance between cell lines of certain species and lineage (e.g. human embryonic kidney cells) and animal cell types (e.g. mouse brain neurons). Therefore, the current process of sensor identification is relatively slow and inefficient, with modest success rate in vivo. Similar to the improvement of CRISPR guide RNA design over the past decade, we envision the application of high-throughput, sequencing-based pipelines to systematically screen RNA sensor libraries that tile the target pre-mRNAs or mRNAs in cell cultures as well as in animal tissues in vivo. These strategies will greatly facilitate the rapid identification of many RNA sensors; they will further generate systematic and quantitative datasets that enable computational and machine learning approaches to derive sensor design principles and prediction algorithms.

#### Sensitivity and dynamic range for sensing cell states

In cell cultures, CellREADR/RADARS/RADAR can detect abundant as well as rare transcripts. Indeed, cellular RNAs expressed in the range of 10-50 TPMs can trigger RNA sensing and payload translation^4–6^. The sensitivity and dynamic range of CellREADR in animal tissues remain to be examined, which requires, a prior, approaches to efficiently identify and evaluate many RNA sensors.

#### Efficacy

Sufficient payload expression level is necessary to achieve intended biological and therapeutic effect. Depending on the delivery method, the efficacy of CellREADR is influenced by generic features such as promoter of expression vector, 5’ and 3’ UTRs, codon usage as well as CellREADR-unique features such as STOP editing (**Figure 4a**). In HEK293 cells with target RNA overexpression, endogenous ADARs can mediate high level payload expression^4–6^. Endogenous HEK293 cellular transcripts can trigger payload expression at lower levels, which can be boosted by co-expressing exogenous ADAR proteins^4–6^. In animal tissues, AAV-delivered constructs yield GFP expression levels that can be readily detected and quantified by immunofluorescence; the expression level may be boosted by incorporating the DART VADAR design to add a positive feedback loop^10^(**Figure 4b**). Binary AAV vectors further allow transcriptional amplification of payload expression, whereby the sensor first triggers the translation of a transcriptional activator tTA, which activates TRE-mediated payload expression in a second vector^4^ (**Figure 4c**). Future improvement of efficacy will optimize each component of CellREADR operation, from readrRNA transcription to RNA stability (e.g. circular RNAs), STOP editing, and translation.

**Figure 4.**
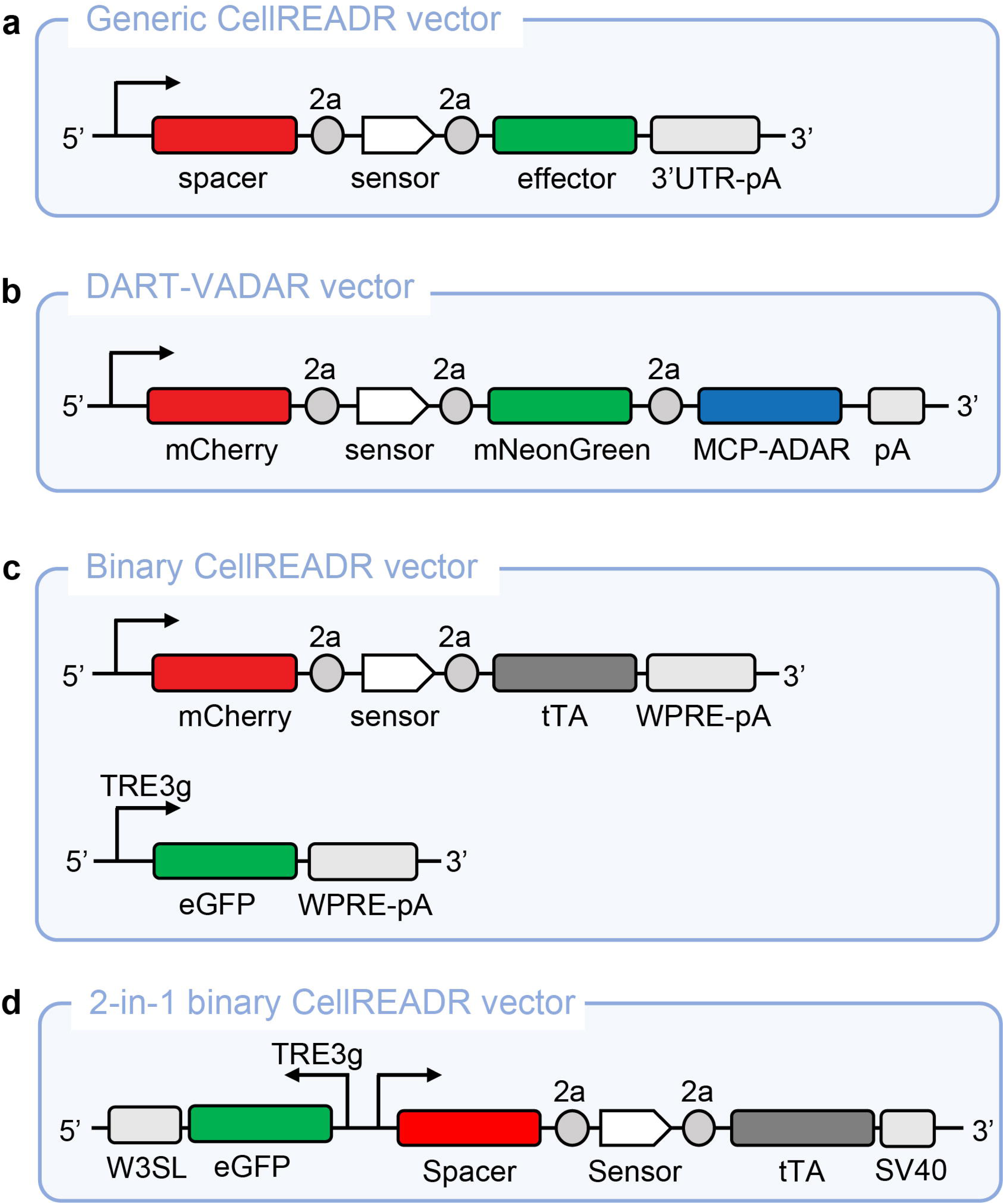
CellREADR expression vectors. **a**, Schematic of a generic CellREADR expression vector. Transcription is driven by an appropriate promoter, which can include an intron. A spacer region is included to reduce STOP read-through of the sensor. The spacer can also be designed to encode a marker protein to indicate transfection/transduction efficiency. The sensor is separated from spacer and effector by sequences coding for self-cleaving peptides (2a). A 3’UTR-polyA region is included for mRNA stability. **b**, The DART-VADAR vector includes a positive feedback loop by adding MCP-ADAR sequence as a second payload and MS2 loops in the sensor region to boost the payload expression level. **c**, With binary CellREADR vectors, RNA sensing first switches on the translation of a transcription activator (tTA) in the *READR* vector, which then amplifies the transcription of an effector controlled by TRE element in the Reporter vector. **d**, 2 in 1 vector is a combination of binary vector and reporter vector in one plasmid to increase the co-transduction rate.

#### Programmability

Cell types and states are often not defined by a single RNA marker but jointly by 2 or more markers. We and others have demonstrated the feasibility of sensing two exogenous RNAs in HEK293 cells^4–6^. The current AND gate design includes two tandemly arranged sensors, each complementary to a different target RNA sequence and contains a separate editable STOP; payload translation is gated by joint sensing and of two target RNAs to remove both STOP codons. This design currently shows relatively low efficiency and likely needs to be improved for in vivo application. In addition to the tandem arrangement of two sensors, it is possible to achieve AND gate through a split payload protein design^4^, whereby the two sensors each separately gates the translation of the N- and C-terminal half of the payload, which then reconstitute following co-expression in the same cells.

Beyond the aforementioned improvements, the ADAR-mediated A-to-I editing machinery presents several constraints on CellREADR. First, recent structure studies demonstrate that ADARs functions as a homodimer^32,33^ and induce editing 35 bp (ADAR1) or 26 bp from structural disruptions (e.g. mismatches)^34,35^; thus optimal ADAR substrates often present as a relatively long stretch of double-strand RNAs of over 50 nucleotides. We and others show that in HEK293 cells CellREADR requires 100 nt or longer substrates to operate. Therefore, small RNAs (miRNAs, shRNAs) are not well suited for RNA sensing by CellREADR. Second, ADAR-mediated A-to-I editing system is absent in plants, fungi, and prokaryotes (RNA editing book); thus CellREADR cannot be directly applied to these organisms. On the other hand, ADAR-based RNA sensor-actuator can be implemented in plant cells by expressing an exogenous human ADAR^P150^ protein^6^. In addition, AAV delivery of is limited by the vector size to 4.7 kb, with a CellREADR payload size up to ∼ 2.5 kb.

### Experimental design

We envision that most investigators will use CellREADR to record and manipulate cells of interest defined by one or more RNA markers. Thus, the experimental design will include the following steps and considerations: 1) RNA sensor design, 2) payload choice and design, 3) readrRNA design and related expression systems, 4) in vitro validation, 5) in vivo delivery, 6) in vivo validation and characterization. CellDEADR implemented through DNA expression vector or in vitro transcribed RNAs differ substantial in experimental design, especially for steps 3)-5). Here we focus on DNA-based CellREADR applications.

#### Sensor design

The locations of the sensor along the target RNA are guided by presence of C**C**A sites for designing editable T**A**G STOP codons in the sensor region (**Figure 5**). The sensor length is generally 200∼300 nt; sensors shorter than 100 nt appear to be less effective^4^. The optimal length remains to be systematically evaluated, especially in vivo. All the in-frame STOP codons in sensor and ATGs after editable ATG need to be mutated, while these mismatches with target may reduce the editing efficiency^4^. Besides, the sensor should not be able to pair with other cellular RNAs in the transcriptome.

**Figure 5.**
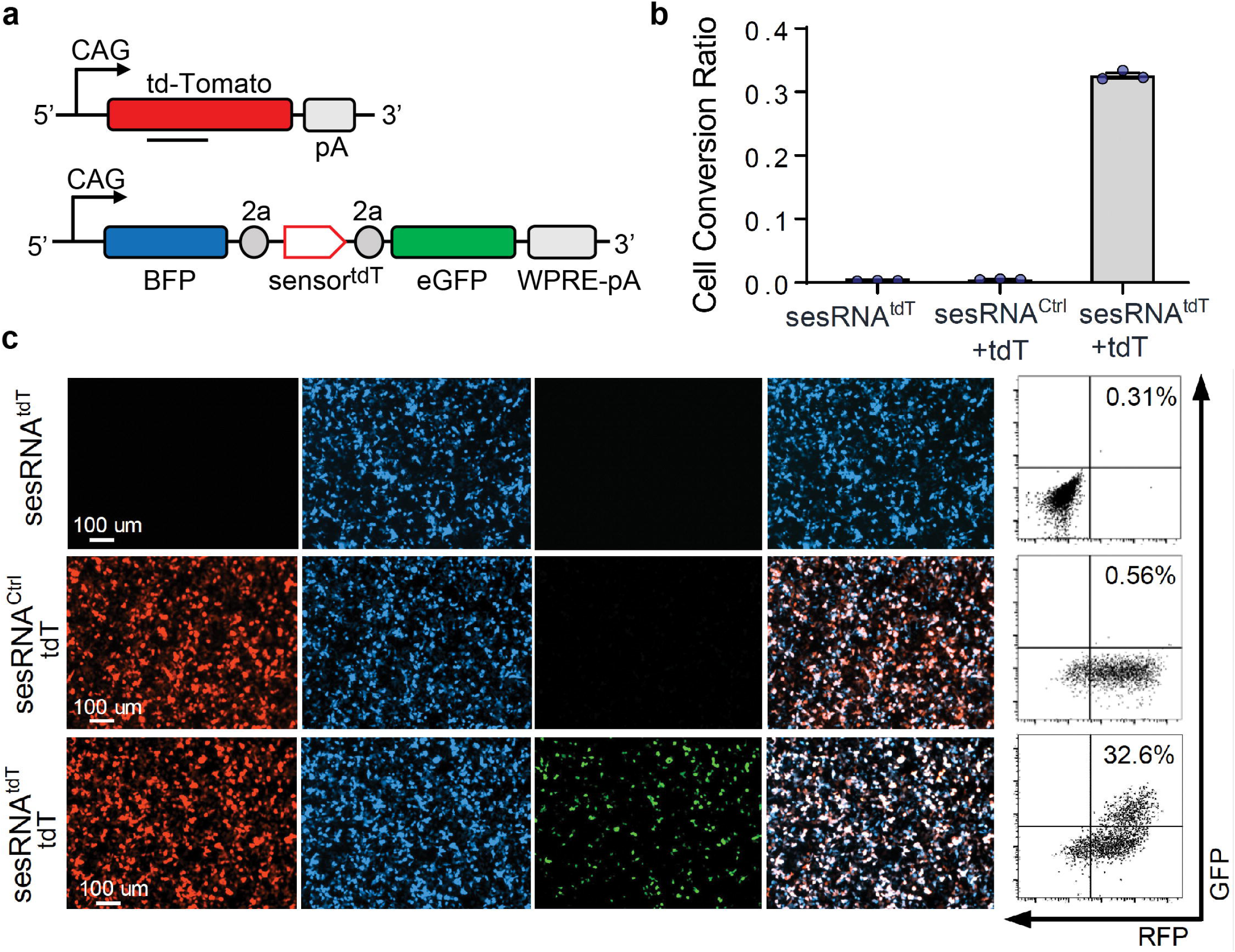
An example of CellREADR in vitro validation. **a**, Schematics of the target RNA vector and CellREADR vector. a, *CAG-tdT* encodes tdTomato target RNA and *READR* ^tdT-GFP^ encodes a readrRNA consisting of a BFP sequence followed by sensor^tdT^ and GFP effector coding sequence. WPRE and 2a are sequences for a virus post-transcriptional regulatory element and a self-cleaving peptide coding sequence, respectively. pA, polyadenylated tail. **b**, Cells transfected with both *CAG-tdT* (tdT) and READR^tdT-GFP^ exhibited robust GFP expression that co-localized with BFP and RFP (bottom). Cells transduced with *READR*^tdT-GFP^ only (top) or *CAG-tdT* and READR^ctrl^ encoding sesRNActrl (middle) exhibited very low GFP expression. Right, FACS analysis of GFP and RFP expression. The percentage of GFP+ cells is indicated. Modified from Qian et al., 2022 with permission. Modified from Qian et al., 2022 Figure1.

Even with these criteria, there are likely many potential sensors distributed throughout the target pre-mRNA or mRNA. In general, we recommend choosing sensors against target regions that are un-disrupted by introns, including single long exons (>200 nt), CDS, 5’ and 3’ UTRs. Sensor against intron sequences also can work well^4^.

Currently we recommend designing 8∼12 sensors for each target to test, depends on the length of the target RNA and the number and locations of CCA anchors in the target sequence.

In the longer term, we envision that large-scale and systematic sensor library screens and characterizations will facilitate computation analysis to derive prediction algorithms for increasingly more reliable and streamlined sensor design.

#### Expression vectors

readrRNAs delivered through DNA expression vectors are expected to undergo the full RNA life cycle as their cellular target RNAs, including transcription, splicing, editing, export, translation, and degradation (**Figure 2**). Thus, expression vectors need be designed to optimize these steps wherever feasible (**Figure 4**).

*Promoters*: The promoter needs to be active in the tissues and cells of interest. Ideally, a “ubiquitous and strong” promoter would serve all purposes. In reality, there are few truly ubiquitous promoters across tissues and species, and some of the more generally active promoters are not the strongest in certain tissues. Thus choosing a well characterized promoter for the tissue and cell types of interest is a necessary first step in vector design.

*Intron*: an intron either in the 5’ or 3’ UTR is often included to mimic endogenously transcribed RNAs and boost expression level.

*Spacer*: this sequence of ∼500 nt or more following the promoter and 5’UTR serves two purposes: 1) to reduce the read-through of STOP codon in the sensor region, 2) to express a marker protein or peptide that can be used to label all the transfected or infected cells.

*Sensor*: as described above.

*Payload*: the size is constrained by the limit of delivery vector. In a typical AAV vector with 4.7 kb overall capacity, the payload coding region can range up to ∼2.5 kb, depending on the size of other components.

*3’UTR and polyA*: This region is important to increase the stability and translation of readrRNA.

Note that a P2A and T2A coding region is included between the spacer and sensor region as well as between sensor and payload sequence. This design ensures that the encoded peptides can self-cleave to produce spacer-encoded marker peptide, a sensor encoded peptide, and the payload protein.

In the *singular vector* described above (**Figure 4a**), the payload translation is directly coupled to the sensor region. To amplify payload expression levels, we have further designed *binary vectors* (**Figure 4c**) in which the sensor first triggers that translation of a transcription activator tTA, which then activates TRE-driven payload expression^4^.

It is possible to package these two components into the same expression vector and fit into the 4.7 kb sized AAV, although the payload size is limited to ∼1.0 kb (**Figure 4d**). This design helps to increase the co-transduction rate.

#### In vitro validation and application

CellREADR can be applied to research in cell lines, iPS, primary cells, and organoids. RNA sensors are validated in cell cultures. DNA expression vectors can be transfected into cells through lipofection or electroporation. Sensor performance can be evaluated and quantified using reporter payloads such as fluorescent proteins or luciferases. For target RNAs that are not expressed in the cell line, a plasmid expressing the exogenous target RNA sequence can be co-transfected with CellREADR vector; and cells without target RNA can serve as negative control for sensor performance. For target RNAs that are endogenously expressed in the cell line, negative controls for sensor testing can be generated by knocking down the target RNA using RNAi^5^or Crispr-mediated transcriptional repression^36^. Sensor performance can be quantified by either fluorescence intensity or conversion ratio over negative control cells (**Figure 5**).

#### In vivo or ex vivo delivery

In vivo delivery can be achieved by viral vectors (AAV, lentivirus, adenovirus) electroporation, and DNA injection into cell nucleus. Among these, AAVs are the most versatile and widely used approach. A key consideration is AAV serotype and tropism: proper capsids need to be chosen and validated for the infection of intended tissues and cells of interest. In addition, AAVs can be delivered either locally by focal injection or systemically by intravenous, retro-orbital, intrathecal, and intraperitoneal injections (**Figure 6**). The choice of proper delivery route is a major determinant of infection regions and efficiency, and thus a key consideration of experimental design.

**Figure 6.**
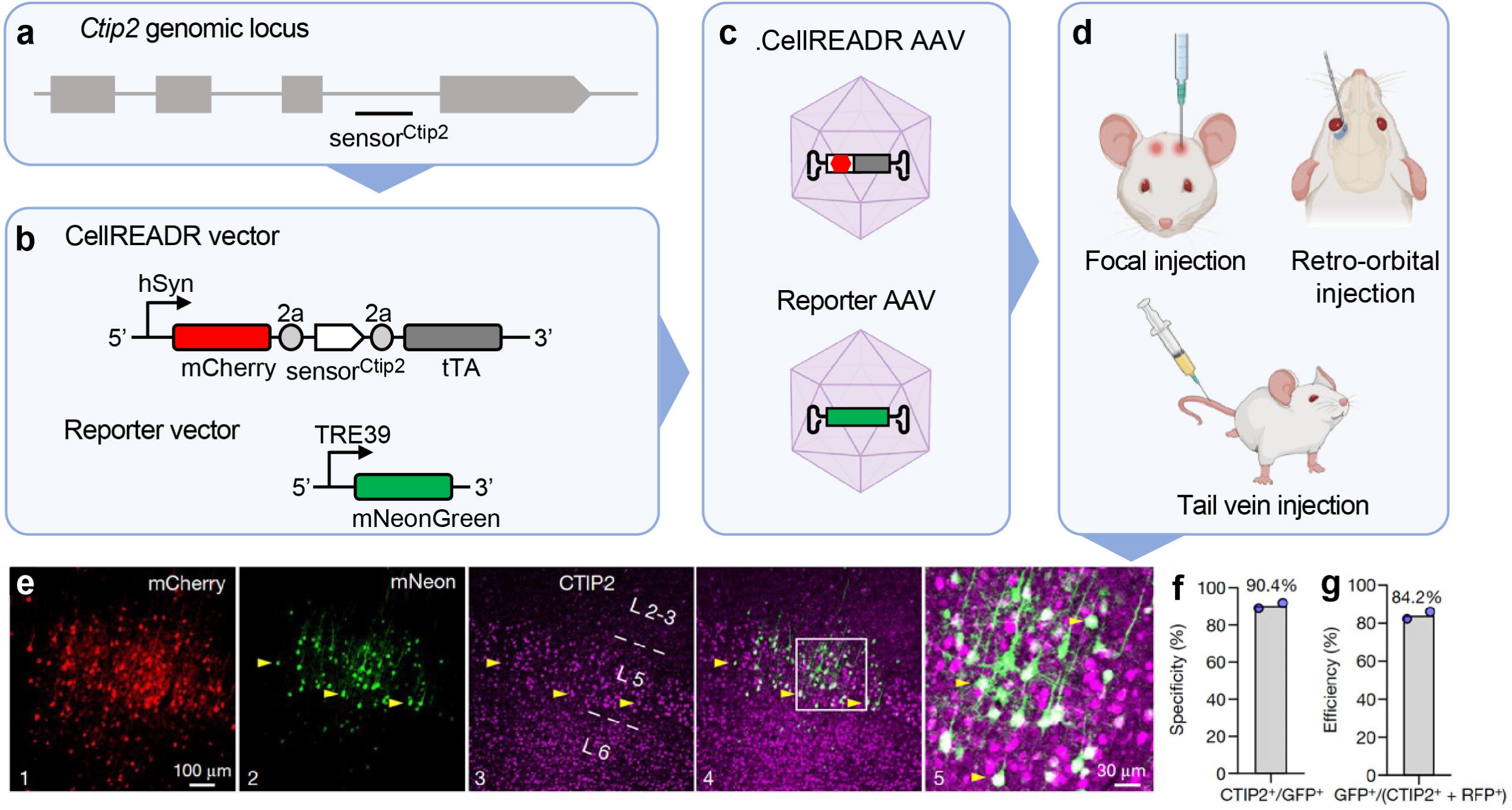
An example of CellREADR in vivo data. **a**, Structure of the mouse Ctip2 gene, showing the location of sesRNACtip2. **b**, Schematic of binary CellREADR AAV vectors for in vivo targeting Ctip2 neurons. mCherry is used as a spacer indicating the transfection. tTA is effector, which can active the expression of mNeonGreen. **c**, CellREADR and reporter are packaged sperately in AAVs with proper serotype. They are mixed before use. **d**, Examples of AAVs delivery routes. **e**, S1 neurons infected with Ctip2 CellREADR AAV expressed mCherry (1); a subset of these in L5b expressed mNeonGreen (2); and their specificity was assessed using CTIP2 immunofluorescence (3–5). The boxed region is expanded in (5). Arrowheads indicate co-labelled cells. **f,g**, Specificity (f) and efficiency (g) of labelling with CellREADR Ctip2binary vectors. Modified from Qian et al., 2022 Figure 4.

**Figure 7.**
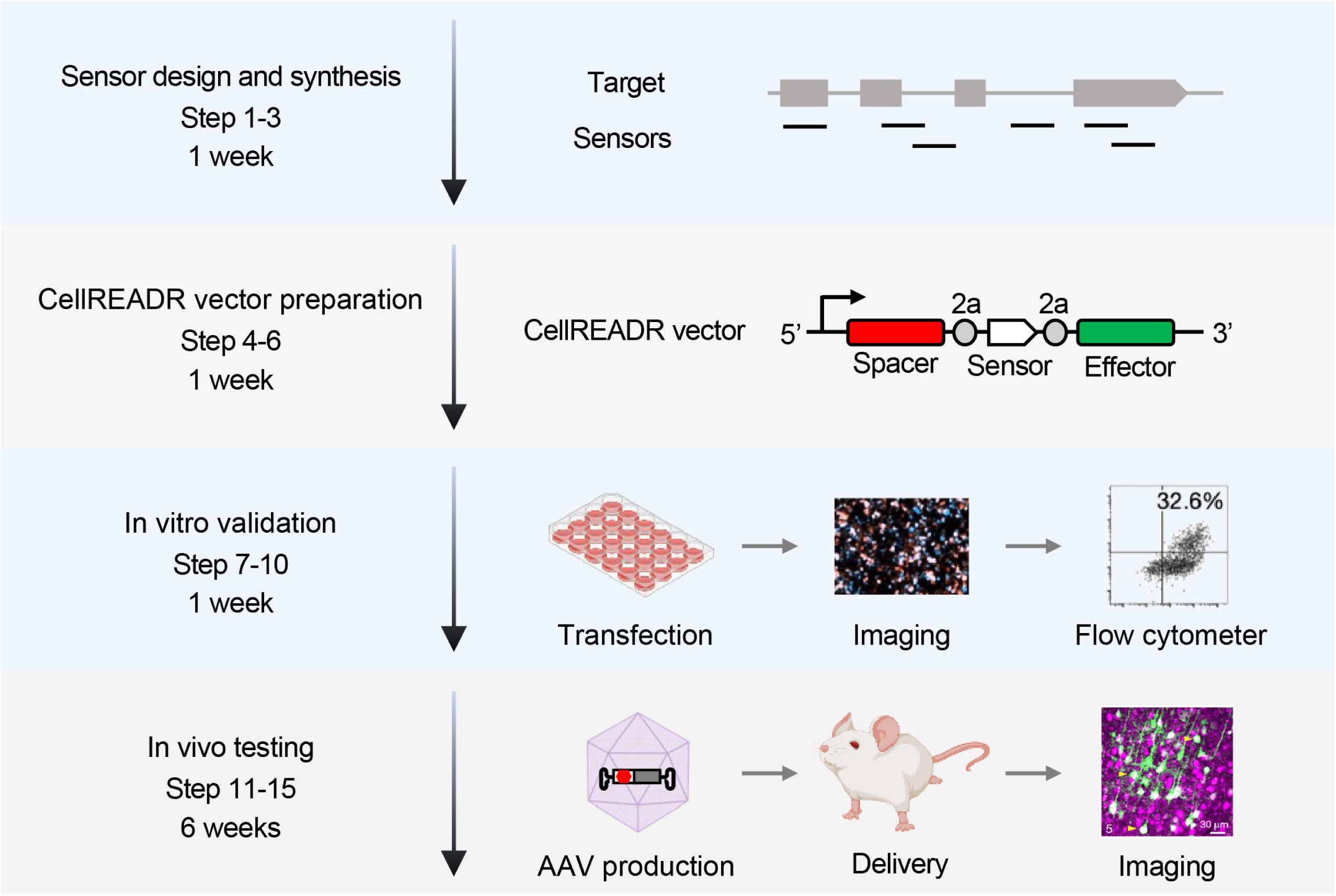
Experimental timeline of CellREADR. Schematic showing the overall workflow and timeline of CellREADR design, in vitro validation and in vivo testing. Overall, the workflow takes 9 weeks to finish.

#### In vivo validation

Usually, 1 or 2 sensors are selected from the in vitro screening and move to in vivo validation. In vivo validation and characterization mostly include three parameters: specificity, efficiency, and efficacy. Typically, using fluorescent proteins (e.g. GFP) as payload is a convenient way to assay and quantify these parameters.

*Specificity* can be quantified as the percentage of payload expressing cells that are positive for target RNA or protein.

*Efficiency* can be quantified as the percentage for target cells that express the payload in the region of interest.

*Efficacy* is the level of payload expression that is sufficient for cell labeling, recording, manipulation.

Here we take the in vivo screening of mouse Ctip2 sensor as an example^4^ (**Figure 6**). The specificity is quantified as the percentage of GFP expressing cells those are positive for Ctip2 immunostaining. The efficiency is quantified as the percentage of GFP expressing cells those are both positive for the spacer mCherry and Ctip2 immunostaining.

When assaying these parameters, it is critical to consider and design proper controls to interpret the outcome. For example, the expression of a marker peptide encoded by the spacer region would indicate that the delivery vehicle and promoter of the expression vector work for the intended tissues and cells. Cells in which target RNA are not expressed or a scrambled “sensor” sequence can serve as negative controls. It is also useful to consider the levels and isoforms of ADAR expression in the cells of interest.

#### Optimizing payload expression

Following the validation of promising sensors, the next step is to modify expression vector and delivery route to optimize payload expression levels and signal to noise ratio (S/N). Inclusion of a minimal version of ADAR as part of payload cassette can boost expression (the DART VADAR design), if the size limitation. Expression cassettes that generate circular RNAs (the tornado vectors) may increase readrRNA half-lift and payload expression. Use of binary vectors can substantially amplify payload expression, but background expression needs to be monitored under an acceptable level. A “2-in-1 binary vector” design combines the READR and reporter components in one plasmid, which can increase the co-transduction rate, but reduces the payload coding capacity for an AAV vector (**Figure 4**).

### Expertise needed to implement the protocol

#### Cell culture

- Molecular biology
- Cell culture
- FACS
- Microscopy

#### Animals

- Viral production
- Animal surgery and injection
- Histology
- Microscopy
- Data analysis and quantification

### Materials

#### Plasmids

- Singular CellREADR vector (Addgene #192063)
- Binary CellREADR vector (Addgene #192066, #192070)
- Reporter vector (Addgene #192064)
- Target vector (Addgene #37825)

#### Plasmid preparation

- Water (Sigma, W4502)
- EcoRI-HF (NEB, R3101)
- AscI (NEB, R0558)
- TopVision Agarose (Thermo Scientific, R0492)
- 50X TAE Buffer (Omega Bio-tek, AC10089)
- 1 Kb Plus DNA Ladder (Invitrogen, 10787026)
- In-Fusion® Snap Assembly Master Mix (Takara, 638948)
- QIAquick Gel Extraction Kit (QIAGEN, 28704)
- QIAprep Spin Miniprep Kit (QIAGEN, 27104)
- NEB Stable Competent E. Coli (NEB, C3040H)
- Carbenicillin (Sigma, C3416)
- LB Agar (Sigma, L2897)
- 2XYT Medium Broth (VWR, 97063-442)
- PureLink™ HiPure Plasmid Filter Maxiprep Kit (Invitrogen, K210017)

#### In vitro screening

- HEK293T cells (ATCC, CRL-3216)
- DMEM (Gibco, 11995073)
- HyClone™ Characterized Fetal Bovine Serum (Cytiva, SH30071.03HI)
- Opti-MEM™ I Reduced Serum Medium (Gibco, 11058021)
- Trypsin-EDTA (Gibco, 25300054)
- PBS, pH 7.4 (Gbico, 10010023)
- Lipofectamine3000 (ThermoFisher, L3000008)

#### In vivo test

- 4% Paraformaldehyde Solution in PBS (Thermo Fisher, J19943.K2)
- PBS (10X), pH 7.4 (Gibico, 70011044)
- OmniPur® Sucrose (MilliporeSigma, 8510)
- Tissue-Tek® O.C.T. Compound (Sakura, 4583)

#### Equipment

- Gel Electrophoresis Systems (e.g. Thermo Fisher Scientific, B2-BP)
- PCR machine (e.g. Thermo Fisher Scientific, A24811)
- Nanodrop (e.g. Thermo Fisher Scientific, 13-400-525) or Qubit (e.g. Thermo Fisher Scientific, Q33327)
- Incubated Shakers (e.g. VWR, 76407-112)
- Centrifuge (e.g. Eppendorf, 4525)
- CO Incubator (e.g. VWR, 10810-902)
- 24-well cell culture plate (e.g. Corning, 3524)
- Fluorescence microscope (e.g. Leica)
- Flow cytometry (e.g. Sony SH800S)
- Anesthesia chamber
- Variable-Speed Peristaltic Pumps (e.g. VWR, 3384)
- Cryostat (e.g. Leica, CM1950)
- Confocal laser microscope (e.g. Leica)

### Procedure

#### Sensor design and synthesis ●TIMING 1 week

1. Select your transcript of interest from NCBI reference sequences. If no specific isoforms/splicing variants are desired, pick the most conserved and highly expressed variant as the target sequence. Ensemble (https://www.ensembl.org/) provides helpful annotations of canonical transcripts. Alternatively, design sensors based on the common regions among the variants.
2. Identify potential sensors using one of the three following options.

A) Web portal

i. Navigate to our web portal located at www.cellreadr.com
ii. Select “Sensors” in menu sidebar.
iii. For a list of potential sensors, use one of the following workflows:

a. Enter target DNA sequence and select compute.
b. For supported species, select species and gene for pre-computed.
B) Manual

i. Generate the reverse complement of your target sequence. Search for tryptophan codons (TGG) in any reading frame of the reverse complementary sequence.
ii. When a tryptophan codon is found, isolate a 201∼300bp region centered on the TGG, and convert the TGG to an editable stop codon TAG.
iii. Locate all STOP codons (TAG, TGA, TAA) in the same reading frame as the editable STOP codon, and convert them to non-STOP codons (e.g. TAC, TCA, TAC). Don’t change T, it will create a mismatched A in targe mRNA, which is not wanted.
iv. Locate all START codons (ATG) downstream of the editable STOP codon in the same reading frame, and convert them to non-START codons (e.g. ATC).
v. Collect regions that meet the following criteria.

a. Contain no more that 5-6 total mismatches with the target sequence.
b. Have no mismatches within 10 bp from the editable STOP codon.
c. Have low sequence similarity with the genome of host organism.
vi. Repeat procedure until entire target sequence had been searched.
vii. Repeat procedure until entire target sequence had been searched.

C) Jupyter Notebook

i. From command line, clone git repository located at https://github.com/zjhuanglab/CellREADR

a. $ git clone https://github.com/zjhuanglab/CellREADR
ii. Run the setup script to initialize the package locally.

a. $ ./setup
iii. Navigate to the folder containing example Jupyter notebooks.

a. $ cd Jupyter-Notebooks
iv. Launch Jupyter Lab.

a. $ jupyter lab
v. Open the template notebook and follow instructions to generate potential sesRNAs.
3. Synthesize the DNA fragments using gBlock/eBlock service from IDT or other companies. In the initial sensor screening, we suggest picking 8∼12 sensors per target that cover different regions of the target transcript. Add 15∼24 bp overlapping sequences on both 5’ and 3’ ends of sensor sequences to match the free ends of digested vector backbones. For most of our Addgene plasmids, we use the following restriction enzymes and overlapping sequences for sensor cloning.

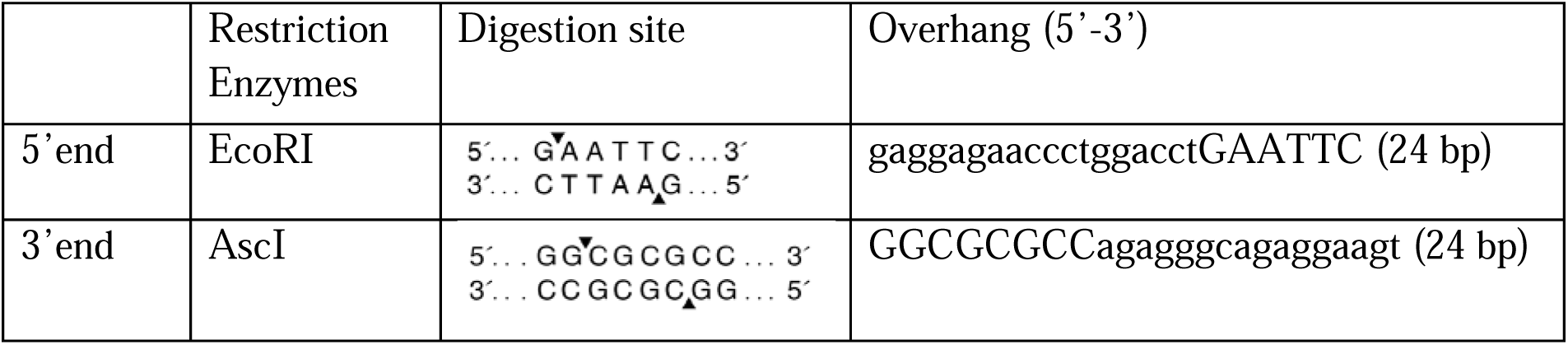

Critical step: make sure the spacer, editable STOP in sensor and effector are in frame before sending to DNA synthesis. Frameshift will result in no effector expression.

#### CellREADR vectors preparation**●**TIMING 1week

4. readrRNA expression vector construction

a. Prepare readrRNA expression vector backbones for in vitro screening (addgene #192063) and choose an appropriate readrRNA expression vector with desired spacer and effector. To start with, we suggest using singular system: use mCherry as spacer and eGFP as effector. Avoid using effector proteins with in-frame ATGs near the 5’ end in the sequence to reduce background effector signal. Digest the vector backbone with EcoRI and AscI for 1 hour at 37 °C. Purify reaction products using gel electrophoresis and QIAquick gel extraction kit following manufacturer protocols.

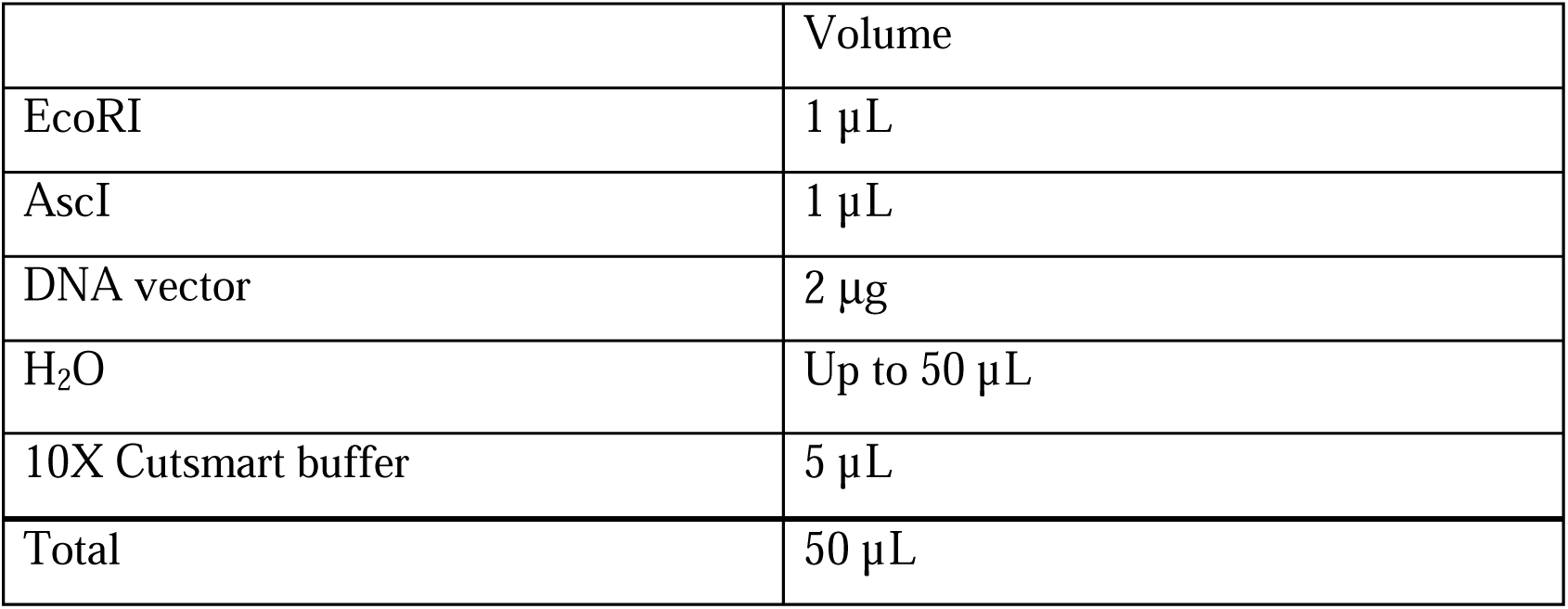
b. Assemble sensor/vector pairs. Mix 3 μL sensor DNA from ITD eBlocks (10 ng/μL), 1 μL vector (∼50 ng) and 1μL 5X Takara In-Fusion® Snap Assembly Master Mix, and then incubate samples in a thermocycler at 50°C for 15 min. Assembled products can be stored on ice for subsequent transformation or at −20°C for long term storage.
c. Bacterial transformation. Thaw competent E. coli on ice. Add 2 μL of assembled product into 10 μL of thawed competent cells and mix gently. Incubate on ice for 15 min then heat shock at 42°C for 30 seconds. Place back on ice and incubate for an additional 1 min. Add 30 μL of room temperature NEB10-beta/Stable Outgrowth Medium to competent cells and incubate at 37°C for 30-60 min in a shaking incubator. Plate cells on a LB agar plate supplied with carbenicillin for selection. Incubate overnight at 37°C.
d. Plasmid purification and Sanger sequence. Pick 1∼2 individual colonies from the plate into 4-5 mL of liquid 2xYT medium supplied with 100 µg/mL carbenicillin. Incubate ∼16 hr or overnight at 37°C in a shaking incubator. Purify the plasmids using QIAprep Spin Miniprep Kit following manufacturer protocols.
5. If the target gene is not or lowly expressed in HEK293T cells, the target RNA expression vector is needed to express the target RNA for in vitro screening. We recommend obtaining the target fragment by PCR, gene synthesis service or commercial plasmids. Clone it into a mammalian overexpression vector (e.g. addgene #37825). Use cells transfected with a non-target RNA expression vector as negative control. **Option**: if it is difficult to get a full long sequence of the desired target, try to split the target sequence into separated constructs.
6. If the target gene is highly expressed in HEK293T cells, siRNA or CRISPRi can be used to perform gene-specific knockdown. Design the knock down vector following the corresponding instructions. Use cells with gene-specific knockdown as negative control. Pause point: All the plasmids can be stored short term at 4°C or long term at −20°C.

#### In vitro screening **●**TIMING 1week

7. Prepare cells for transfection. Seed HEK293T cells in a 24-well plate at 1×10^5^ cells per well the day before transfection (day 0). Culture the cells with complete DMEM media at 37°C in 5% CO_2_ incubator.
8. On day 1, transfect cells using Lipofectamine3000 following manufacturer protocols. For sensors targeting exogenous genes, we routinely co-transfect 300 ng readrRNA expression vector and 100 ng corresponding target RNA expression vector. For negative control, replace the target RNA expression vector with 100 ng non-target plasmid. For sensors targeting endogenous genes in HEK293T cells, a higher amount of singular readrRNA expression vector (e.g. 500 ng per well) can be used. Incubate cells at 37°C in 5% CO_2_ incubator for 2∼3 days.

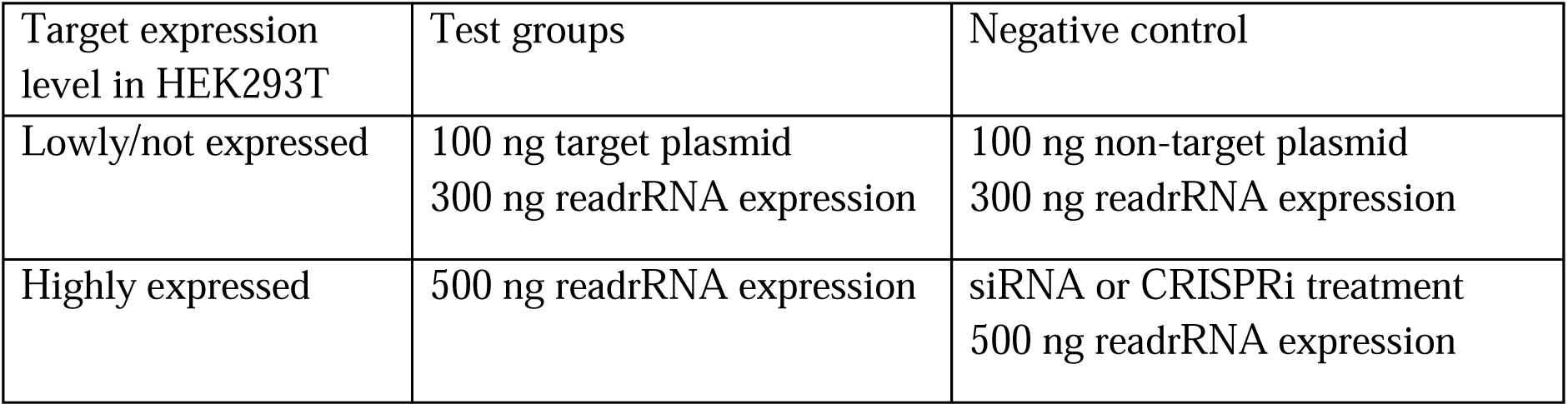
9. On day 3∼4, image cells under fluorescent microscope to get an estimate of sensor performance. The percentage of the spacer signal (i.e. mCherry signal if used as spacer) in all cells reflect the transfection rate. When transfection rate is similar among samples, the fluorescence intensity of effector (i.e. eGFP signal if used as effector) can be used to assess sensor performance. A specific and efficient sensor should show both high effector signal when target RNA is present and low background signal when target RNA is absent.
10. Quantify sensor efficiency with flow cytometry. Each well of transfected cells is dissociated with 100 μL 0.05% Trypsin and collected in a 1.5 mL tube with 500 μL complete DMEM media. Centrifuge the cells at 300*g* for 3 min, aspirate supernatant and resuspend cells in 200 μL 1xPBS. Keep cells in the dark and on ice during the analysis. Run the flow cytometry analysis with proper single fluorophore controls for compensation, depending on the choices of fluorescent proteins for spacer and effector. Collect the data and analyze it with FlowJo v10. The sensor efficiency is quantified by cell conversion rate, defined by effector expressing cells (effector+) among all the transfected cells expressing spacer signal (spacer+), i.e. [effector+cell] / [spacer+ cells]. Alternatively, median fluorescent intensity can be used to quantify effector signal strength of transfected cells.

#### In vivo test **●**TIMING 6 weeks

11. Prepare AAV plasmids for in vivo testing.

a. readrRNA expression vector. Choose an appropriate AAV readrRNA expression backbone with desired promoter, spacer and effector. As a general guidance, the chosen promoter should drive high and stable expression of CellREADR in the desired cell population, regardless of the inherent cell-type specificity of the promoter. The spacer should provide reliable transfection control in the testing condition, and the effector should allow desired manipulation of the target cell type. Avoid using effector proteins with in-frame ATGs near the 5’ end in the sequence. In the case of mouse Ctip2 sensor tested in vivo, the human Synapsin promoter was used to ensure high readrRNA expression in neurons, mCherry was used as spacer to indicate successful AAV transduction, and tTA was used to amplify mNeonGreen signal by activating TRE3g promoter (Figure 6b). Select the best sensor with high cell conversion rate and low background from in vitro validation, and assemble it into the AAV backbone following the cloning method described above. **Option:** positive and negative controls. To create a positive control sensor, mutate the designed TAG STOP codon in the selected sensor back to TGG, and assemble it to the same readrRNA expression backbone. This positive control sensor should express effector constitutively in all transduced cells, allowing users to evaluate the choice of promoter, AAV serotype and injection method. To create a negative control sensor, a scrambled sequence based on the selected sensor, can be made without introducing an editable stop codon. This negative control sensor cannot express its effector in any cell.
b. Reporter vector. When using binary CellREADR vectors, choose a desired effector and clone it in pAAV-TRE3g plasmid (addgene #192064) with SalI and SpeI.
c. Confirm AAV plasmid sequence by whole-plasmid sequencing, and make sure the plasmid includes two intact ITRs. When a reporter AAV vector is needed, confirm the whole sequence too. Prepare plasmid DNA using PureLink™ HiPure Plasmid Filter Maxiprep Kit following the manufacturer protocols. Pause: All the plasmids can be stored short term at 4°C or long term at −20°C.
12. AAV packaging. Choose a proper serotype according to the target cell types, organs, animal species and delivery methods. Package CellREADR AAV and reporter AAV separately following the AAV production and purification protocols from Addgene (https://www.addgene.org/protocols/aav-production-hek293-cells/; https://www.addgene.org/protocols/aav-purification-iodixanol-gradient-ultracentrifugation/). Titration of AAVs can be performed by qPCR using SYBR Green technology following the protocol from Addgene (https://www.addgene.org/protocols/aav-titration-qpcr-using-sybr-green-technology/). In the case of mouse Ctip2 sensor tested in vivo, Ctip2 readrRNA plasmid and reporter plasmid were packaged with AAV-DJ serotype and injected into mouse cortex by focal injection (Figure 6c,d). Pause point: AAVs can be stored at 4 °C for short term (2 weeks), or −80 °C for long term.
13. AAV delivery and incubation. CellREADR AAVs can be delivered through focal injection, tail vein injection, retro-orbital injection, or intraperitoneal injection. The dose of AAV for injection varies depending on the type of injection and the animals being injected. If binary vectors are used, mix CellREADR AAV and reporter AAV before injection. Typically, after injection, we will incubate CellREADR in animals for 3∼4 weeks. **Critical step**: carefully select an appropriate dose of AAV, injection methods and AAV serotypes. Optimize these parameters to ensure sufficient sensor expression. Failed injection or insufficient dose will result in no or low expression.
14. Tissue collection and processing. After AAV incubation, anesthetize the injected mouse and perform transcardial perfusion with 1xPBS followed by 4% paraformaldehyde (PFA). Dissect the mouse and collect desired organs. Place the organs in 4% PFA solution overnight, and then dehydrate and embed them using preferred reagents. Section tissue using preferred technique. For mouse brain, we use 30% sucrose in PBS to dehydrate and embed the tissue in OCT. Freeze the tissue and section with cryostat. Collect the sections in PBS and store at 4°C for imaging and staining. Pause point: sections can be store in PBS at 4°C for short time. Critical step: For best tissue quality, perform dissection quickly. Heart should still be beating when needle is inserted to start perfusion. Should observe body twitching and tail flicking when perfusing with PFA. Body should be stiff following perfusion.
15. Imaging and quantification. Cellular target mRNA or protein can be stained by mRNA in situ hybridization or immunostaining, respectively, following appropriate protocols. AAV-transduced cells (expressing a marker peptide encoded in the spacer region) and payload protein can be stained and quantified by immunostaining. Obtain confocal images using appropriate filters and settings. Specificity is calculated as the percentage of effector-expressing cells that are positive for target RNA or protein marker ([target+ cells] / [effector+cells]. Efficiency is calculated as the percentage of target cells that express the effector protein ([effector+cells] / [spacer+ target+ cells]). In the case of mouse Ctip2 sensor tested in vivo, the target RNA expression was confirmed by immunostaining of Ctip2 antibody. Successful AAV transduction was indicated by mCherrry expression. The sensor payload tTA was expressed to transcriptionally activate TRE3g promoter, leading to mNeongreen expression (Figure 6e). The specificity of Ctip2 sensor was quantified as Ctip2+ cells/GFP+ cells. The efficiency was quantified as GFP+ cells/(RFP+Ctip2+ cells) (Figure 6f,g)

Timing:

Step 1-3: Sensor design and synthesis, 1 day

Step 4-6: CellREADR vectors preparation, 1 week

Step 7-10: In vitro screening, 1 week

Step 11-15: In vivo test, 6 weeks (including 2 weeks for AAV packaging)

Troubleshooting:

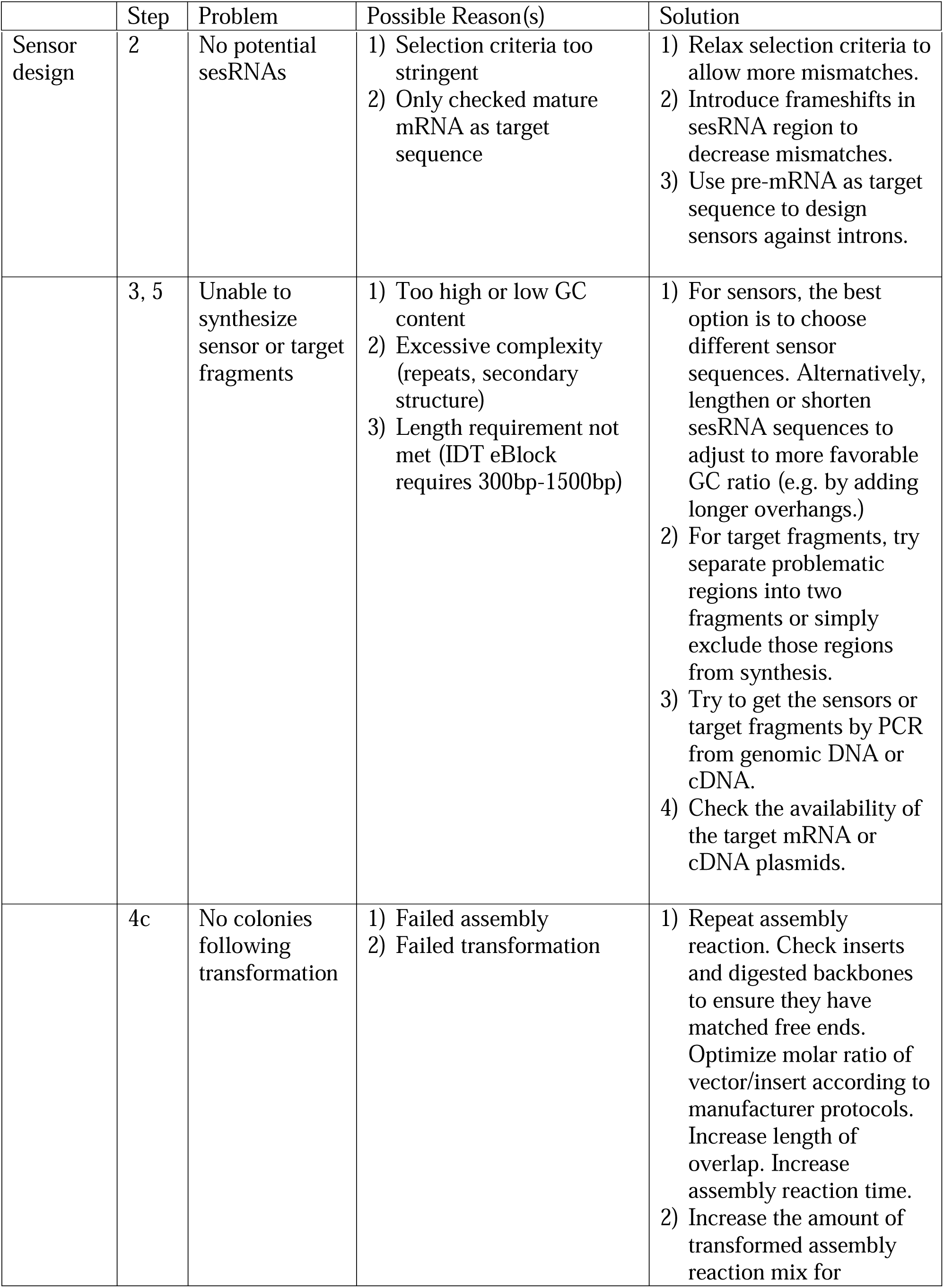
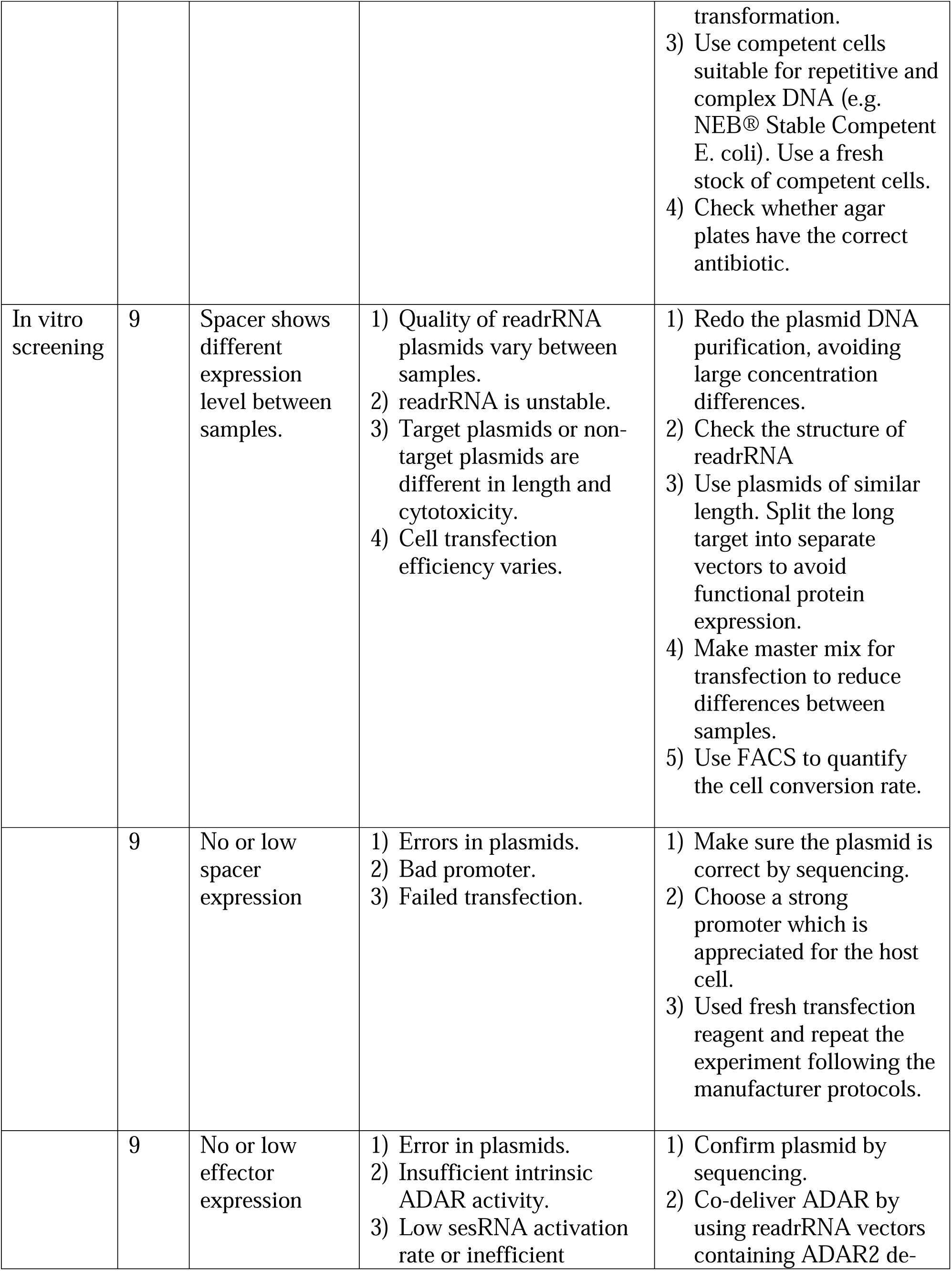
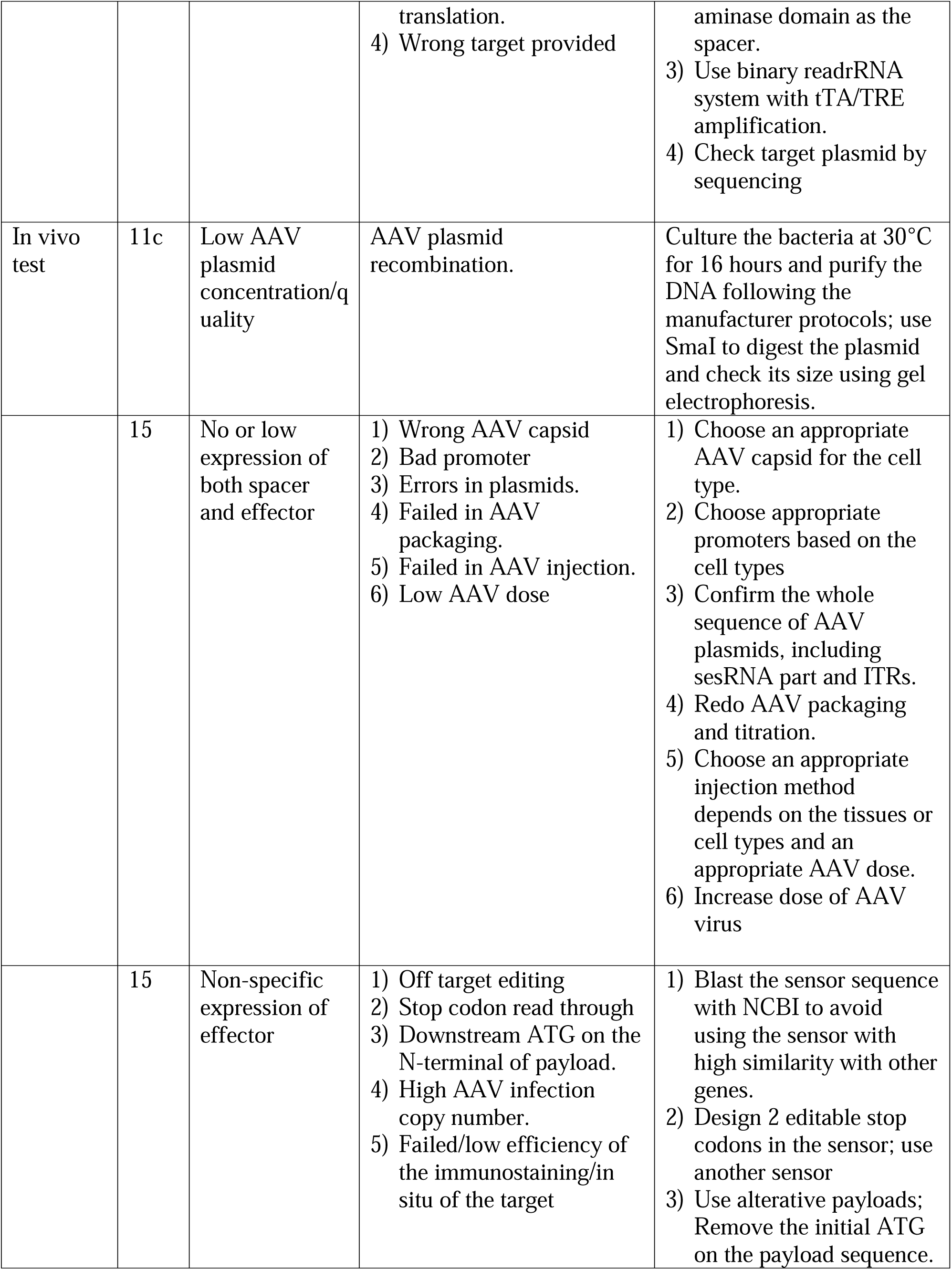
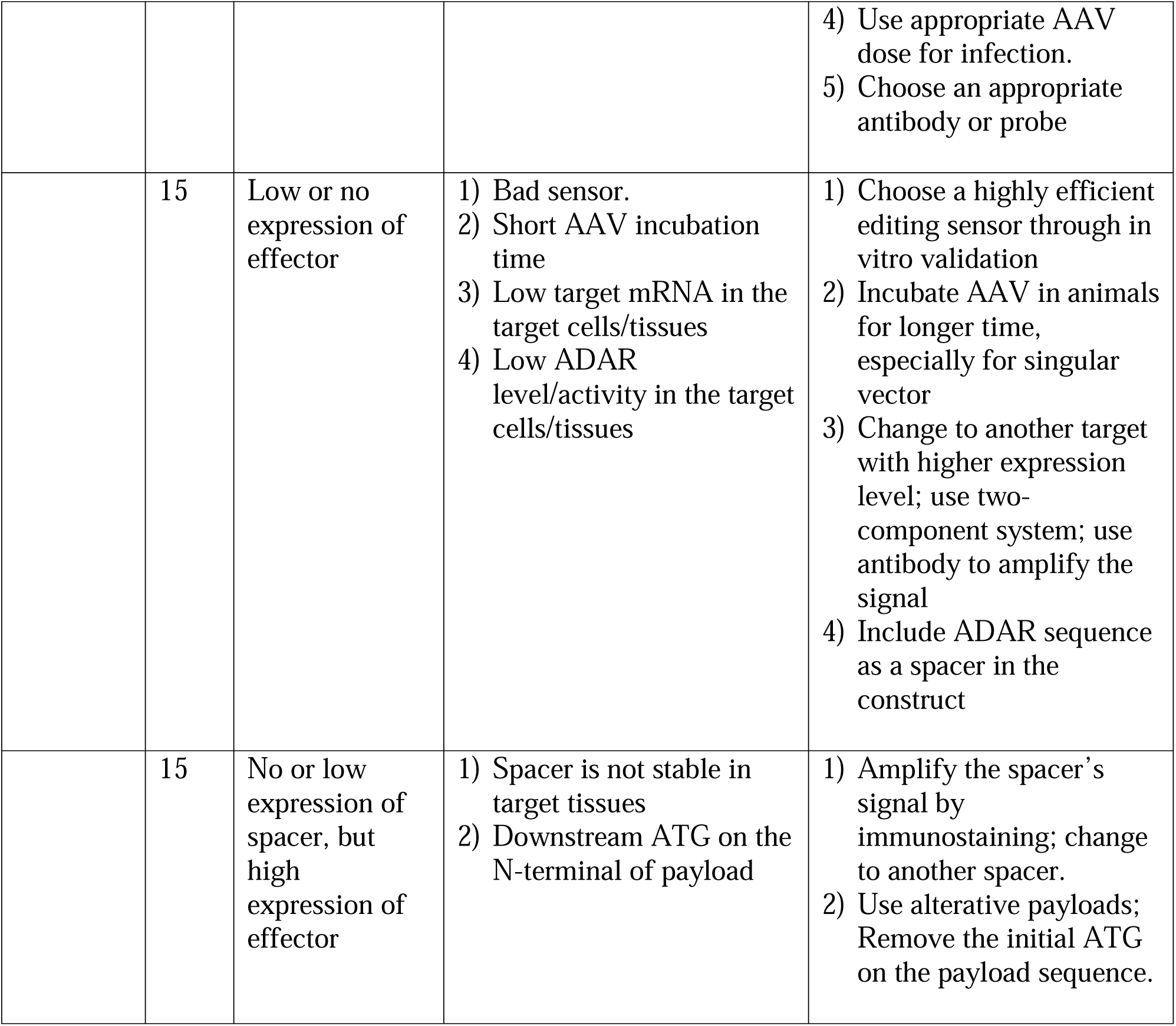

### Anticipated results

Based on the target sequence, sensors can be designed and assembled into CellREADR vectors. By screening 8 ∼12 sensors in vitro, we expect to select at least 2∼3 highly-active sensors with high cell conversion rate and low background. These selected sensors can be used for further in vivo validation.

Singular or binary CellREADR vectors can be used for in vivo testing. When singular vectors cannot express effectors at a desired level, binary vectors can amplify effector signal with tTA-TRE transcriptional amplification system. A successful readrRNA targeting gives high specificity and efficiency.

After selecting the best-performing sensor from the in vivo testing, the fluorescent protein effector can be replaced with a functional protein to specifically manipulate the targeted cell population.

## Acknowledgements

This work was supported in part by NIMH grants 1DP1MH129954-01to Z.J.H. and NIH grants R01MH113005 to J.H., J.G. and Z.J.H. We thank W. Zhong for helping with processing of Figure 3.

## Contributions

Z.J.H, X.Y., K.W., wrote the introduction and protocol. J.H. J.G. contributed to web portal design. S.Z., X. G., Y.Q., contributed to trouble shooting sections. All authors edited the manuscript.

## Competing Interests

Z.J.H. and Y.Q. have filed patent applications on CellREADR. Z.J.H is a co-founder of Doppler Bio. All other authors have no conflicts of interest to declare.

